# Theory and application of an improved species richness estimator

**DOI:** 10.1101/2022.05.02.490342

**Authors:** E. W. Tekwa, Matthew A. Whalen, Patrick T. Martone, Mary I. O’Connor

**Affiliations:** Department of Zoology, University of British Columbia, Vancouver, BC, Canada; Hakai Institute, Heriot Bay, BC, Canada; Department of Botany, University of British Columbia, Vancouver, BC, Canada

**Keywords:** species richness, biodiversity, bias, occupancy, clustering, bootstrapping

## Abstract

Species richness is an essential biodiversity variable indicative of ecosystem states and rates of invasion, speciation, and extinction both contemporarily and in fossil records. However, limited sampling effort and spatial aggregation of organisms mean that biodiversity surveys rarely observe every species in the survey area, which introduces bias to the estimated richness and inaccuracy to comparisons of communities across space and time. Here we present a nonparametric, asymptotic, and bias-corrected richness estimator, *Ω*_T_, by modelling how spatial abundance characteristics affect observation of species richness. We conduct simulation tests and applied *Ω*_T_ to a tree census and a seaweed survey. *Ω*_T_ consistently outperforms common estimators in balancing bias, precision, and difference detection accuracy. Our results provide theoretical insights into how natural and observer-induced variation affects species observation and support *Ω*_T_ as a promising and application-ready richness estimator for a wide variety of data.

## Introduction

A central objective of biodiversity science is to understand how species richness changes across space and time [1]. Not only is species richness important for today’s ecosystem and human health assessments [2–4], it is essential to inferring paleoecological patterns and mass extinction events [5]. Species richness estimation techniques are among the most highly cited works in ecology [6–8]. Further, estimating missing species or classes from limited observations is a fundamental problem in other fields, including genomics [9], cryptography, machine learning, and linguistics [10].

While it is clear that global extinction rate exceeds baseline levels [11,12], synthesis of species richness timeseries have suggested both gain and loss of local species richness at local scales [13–16]. Observed species richness and detection of differences across communities or time are prone to data bias [17] and undocumented errors resulting from varying survey and sampling effort, design, and observer skill [18]. These problems are more acute for species richness than for other biodiversity metrics [19], in part because species richness is sensitive to detection of rare species. Data-intensive methods such as multi-species occupancy models can estimate richness of targeted species groups, but they require repeated field surveys and are sensitive to models of abiotic and biotic drivers [20,21]. For many general applications and meta-analyses, it is desirable to have an asymptotic (finite and effort-independent) richness estimator that can both correct bias (uncover true absolute richness) and accurately detect spatial and temporal differences across communities.

In most places, the best biodiversity data we can hope for are individual counts identified to the species level. One obvious way to improve biodiversity estimation is to increase sampling effort, but this is costly and not possible for some taxa, habitats, and historical samples. Insufficient sampling effort can reduce diversity estimation precision and undermine efforts to detect change over time, but a robust, unbiased estimator can recover missing observations to some extent. The most robust asymptotic richness estimator that uses abundance data remains the Chao1 method [6,22]. Chao1 and related estimators are based on Turing’s sampling theory that relates the number of unobserved classes (codes or species) to observed rarity [22,23]. These methods are expected to be inaccurate with spatial heterogeneity, low sampling effort, and heterogeneous observation probabilities. The latter issue has been tackled using models requiring tuning, but these methods can be unstable and still under development [24]. Further, bias to both estimates and confidence bounds remain intractable problems [25]. There is a need to develop new statistical methods that fully utilize information in common survey data, without having to separately estimate additional quantities that may be less reliable than raw richness itself [20].

Here we outline five technical problems that a new asymptotic estimator should account for. The first two are human-induced obstacles to species observation: low fraction of individuals observed and low number of patches (sites, transects, or quadrats) sampled. The last three are natural properties of biological spatial distributions that also reduce the number of species observed: low abundance, patch occupancy, and spatial clustering within patch (Figure 1). These variables will be precisely defined, and we use them to mechanistically derive observation probabilities and a set of what we will call *Ω* bias-corrected asymptotic richness estimators. This approach departs from both Turing’s reliance on rarity, which is prone to noisy observations, and from earlier parametric extrapolations that have weak justifications [22]. Using the derivations, we highlight how distributions of the variables affect richness observations. Next, we evaluate the performance of two proposed *Ω*-estimators relative to five other best-established asymptotic estimators (Chao1, Chao2, ACE, Jackknife1-abundance, Jackknife2-incidene) using simulations designed to challenge each estimator through the five problems introduced above. Six statistics measure bias (under- or over-estimation tendency), precision (repeatability of estimates), and accuracy (error magnitude and spatial/temporal trend identification). We additionally devise an improved bootstrapping procedure for confidence bounds that account for the spatial- and species-dependence between individuals. To test our method with empirical applications, we used two multi-year, multi-patch datasets: the Barro Colorado Island tree census [26] and the British Columbia intertidal seaweed survey [27], where the probable true temporal trends are, respectively, decreasing and not changing. We conduct spatial subsampling and local downsampling experiments on the real datasets to explore how much different estimators can recover absolute richness and detect temporal trends. Finally, we create Matlab and R functions for estimating richness via a suite of asymptotic methods.

**Figure 1.**
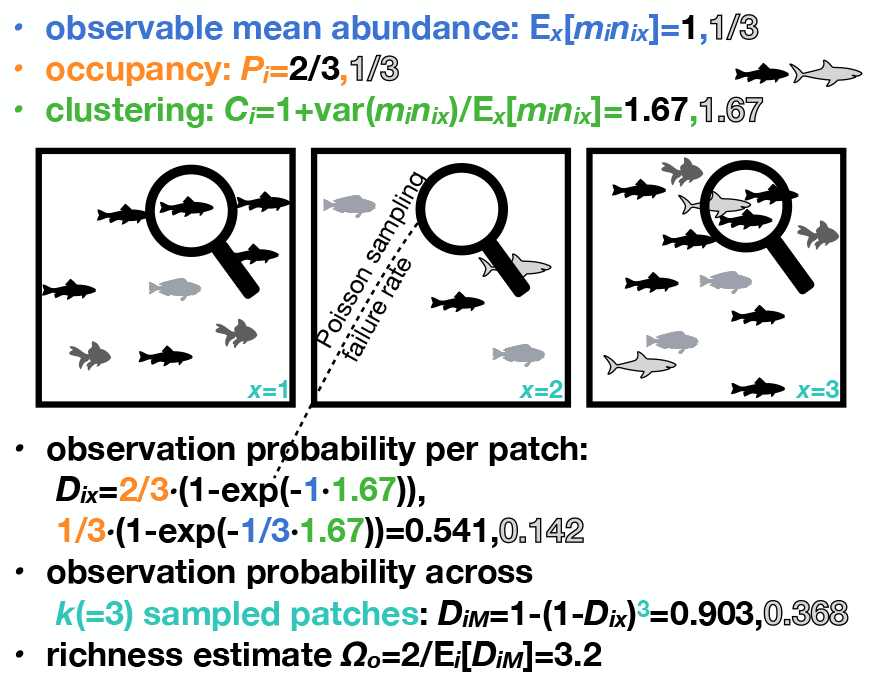
From spatial statistics to richness estimate. Spatial statistics are used to infer observation probabilities, leading to a bias-corrected richness estimate *Ω*_*o*_. The statistics are observable mean abundance *mn* (blue), occupancy *P* (orange), clustering *C* (green), and sites sampled *k* (cyan) from the two observed species (black and outlined grey numbers).

## Methods

### Richness Estimator Derivation

Here we define variables that can be measured from a spatially partitioned abundance dataset, with the aim of arriving at a bias-corrected richness estimator. Our strategy involves the following steps (Figure 1B). We begin at the finest spatial resolution sampled (“patches”) where individuals are counted, across which we track how both spatial heterogeneities and observation errors (through spatial abundance statistics) affect the probability that a species is observed at least once at the larger community level across patches. We then use spatial abundance statistics of observed species to estimate true richness in the underlying community, which is the observed richness divided by some expected observation probability.

In a community defined by patches (indexed *x*) belonging to the spatial set *M*, each species *i* has a true mean density *n*_*i*_, which is the total number of individuals at scale *M* divided by the number of patches *X*, or equivalently the individuals per patch in *k*_*i*_ randomly sampled patches. The fraction of individuals observed at *x, m*_*ix*_, is also understood as the individual catchability of species *i* [28]. Thus, *m*_*ix*_*n*_*ix*_ is the local “observable abundance”, which will serve as the first (non-spatial) random state variable in our subsequently developed richness estimator.

In contrast to the above community scale, let us descend to the local patch scale where observation occurs. At each local patch (“observation neighbourhood”) where an individual of species *i* occurs, there tends to be *n*_*i:i*_ individuals (counting the focal individual itself) on average because of spatial variance [29]. Note “on average” here means per individual, not per occupied patch, the latter being equal to or smaller than the former because some patches have more than one individual. The two subscript *i*’s indicate that *n*_*i:i*_ is the local density of species *i* from the perspective of an individual of species *i*, which contrasts from the global density *n*_*i*_. The quantity *n*_*i:i*_ can be understood as the Lagrangian description of spatial distribution (i.e. tracking other individuals from a focal individual’s perspective), which can be precisely translated to abundance at patch *x*: *n*_*ix*_, the Eulerian description (i.e. tracking individuals from a fixed spatial perspective) in fluid dynamic terminologies that gave rise to modern spatial ecology [30]. The spatial variance *var*_*x*_(*n*_*ix*_) is the variance in the true number of individuals of species *i* across all patches *x*. It can be shown that the mean local density *n*_*i:i*_ is a function of spatial variance, with variance having a normalization factor *k*_*i*_, the number of patches sampled, rather than the usual 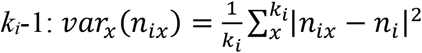 [31,32].

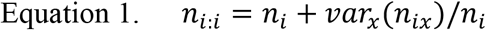

This local density is the true expected local abundance that the human observer samples from in an occupied patch, because in such a patch the observer takes the same perspective as an individual of species *i*. We define the first spatial heterogeneity quantity that will serve as our second random state variable – the within-species clustering across space *C*_*i*_ – as [29,32]:

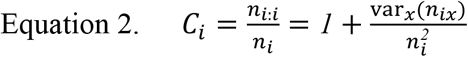

Note the following identity allows us to simplify Equation 2 in terms of the observable abundance *m*_*ix*_*n*_*ix*_ across patches *x* instead of the generally unobserved true abundance *n*_*ix*_, if the fraction of individuals observed does not differ across patches (*m*_*ix=*_*m*_*i*_) – an assumption we will carry forward:

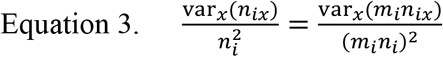

The expected local observable abundance in an occupied patch is thus *m*_*i*_*n*_*i:i*_ =*C*_*i*_*m*_*i*_*n*_*i*_. Using this mean the local observation process can be described by a non-spatial Poisson sampling process (sampling with replacement). This local Poisson sampling process should not be confused with a Poisson spatial distribution across patches, which was not assumed. Let the local observation probability of species *i* at an occupied patch (the spatial set *X*_*io*_) be 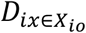. This probability is one minus the rate of failure to observe:

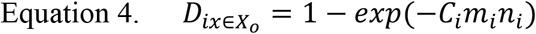

Now we zoom back out to the community level *M* and consider spatial processes across multiple patches that may be occupied or not. The second spatial heterogeneity quantity – our third state variable – relevant to this level is occupancy *P*_*i*_, which is the number of occupied patches *k*_*oi*_ by species *i* divided by *k*_*i*_:

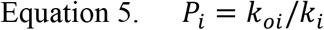

Occupancy is related to clustering through spatial variance, but this relationship is mediated by true mean abundance [33], which we generally cannot observe from biodiversity surveys because of imperfect observations. Thus, we treat occupancy and clustering as separate dynamic variables but, as we will see, our statistical method accounts for the effect of their covariance. The observation probability of species *i* per random patch *x* is the occupancy (Equation 5) times the conditional local observation probability (Equation 4) when the patch is occupied.

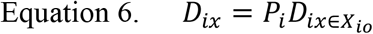

The observation probability of species *i* within a community *M* is one minus the cumulative probability of failure to observe, the latter being the probability of failure to observe within patch raised to the power *k*_*i*_ (our fourth state variable) if patches are randomly picked. The observation probability at scale *M* (*D*_*iM*_) is:

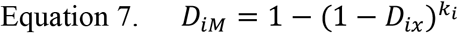

If patches are not randomly picked, then *k*_*i*_ should be the effective sample size. As an extreme example where only one patch is repeatedly sampled at the same time in exactly the same way, effectively *k*_*i*_=1. In practice however, even sampling the same patch repeatedly would generate different individual and species counts when catchability *m* is below 1 and/or individuals move between patches during the finite time separating samples. Ignoring the non-independence of spatial samples would inflate *D*_*iM*_, but defining what is meant by effective sample size will require nuanced considerations of how the spatial sampling design relates to the community’s spatial distribution. We will assume *k*_*i*_ samples are independent and leave effective sample size as a venue for future improvements.

The raw observed species richness *S* can be written without additional assumptions as the true richness *Ω* multiplied by the expectation of *D*_*iM*_ across truly present species *i* as a function of ***Φ***, which includes *R* state variables *ϕ*_*r*_. In our observation process model, *R*=4 and *ϕ*_*1*_ to *ϕ*_*4*_ are *m*_*i*_*n*_*i*_, *C*_*i*_, *P*_*i*_, and *k*_*i*_.

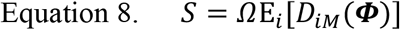

We can stop here and define *Ω*_o_ (Equation *9*) as our first proposed bias-corrected richness estimator using the average observation probabilities based on state variables 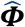 measured from the set *S*_*o*_ of observed (rather than truly present) species (see Figure 1 for numerical example). All quantities with a 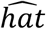 denote observed rather than true values in the subset *S*_*o*_. The estimator *Ω*_o_ contains survivorship bias [34] – a type of sampling bias stemming from the difference between 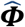 and ***Φ***, because 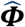 is only measurable for observed species. For intuition, observed species will likely have higher abundances than the average of truly present species because rare species tend to be missed.

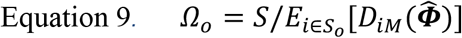

An alternative is to write E_*i*_[*D*_*iM*_(***Φ***)] in Equation 8 in terms of a second-order Taylor expansion (with *r* and *q* being variable indices), which provides insights into how means and variances in different statistics affect the observation process:

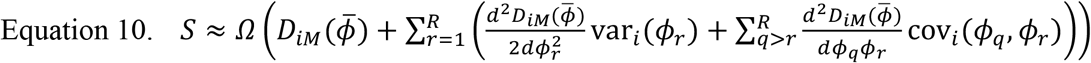

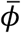 stands for the true average values of the *R* state variables across truly present species. If there are observation errors, higher moments are increasingly harder to estimate and may destabilize the estimator. We thus propose an idealized estimate for *Ω*, the second-order Taylor approximated richness *Ω*_*TC*_, with *TC* indicating that the spatial abundance statistics are the “corrected” or unbiased statistics for all truly present species and not just observed species:

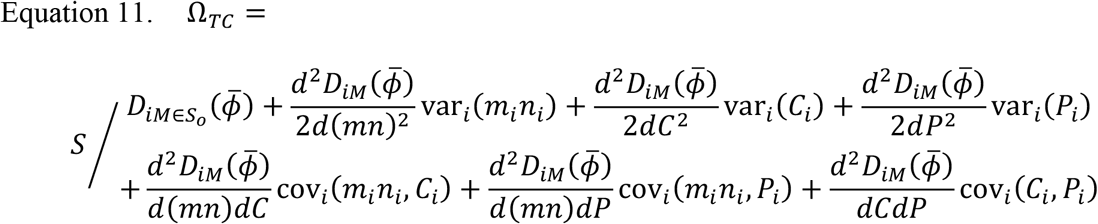

*Ω*_*TC*_ contains a different set of bias than *Ω*_o_. The potential bias for *Ω*_*TC*_ comes from the Taylor moment truncation beyond the second order terms but contains no survivorship bias (the spatial statistics are from ***Φ***). However, *Ω*_*TC*_ cannot be implemented in real data because we do not know ***Φ*** containing 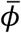 and *ϕ*_*r*_ across all truly present species. We thus propose a Taylor approximated richness *Ω*_T_ that uses directly measured variables 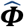 in place of ***Φ*** in Equation 11, which is operational on real biodiversity survey datasets but contains survivorship bias. This estimator is:

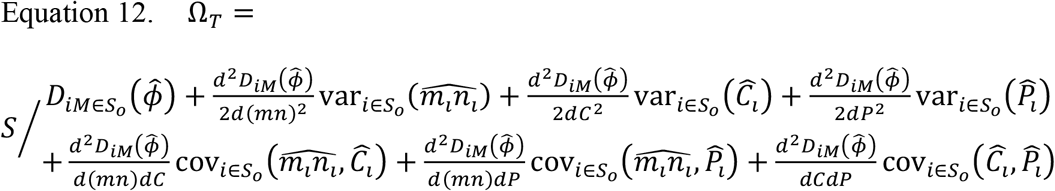

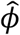 stands for the observed across-species averaged value set 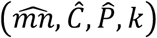 and 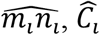, and 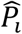 are the observed values for each species *i*. We assumed that *k* is always known and identical for all species in a survey, hence there is not a second-order correction term associated with variance in *k*. We define the seven terms in the denominator of Equation 12 as “correction factors”: 0^th^-order, *V*_*mn*_, *V*_*C*_, *V*_*P*_, *C*_*mn,C*_, *C*_*mn,P*_, and *C*_*C,P*_. *Ω*_*T*_ looks complicated but is computationally trivial and effectively takes no more time or data than *Ω*_o_ or other abundance-based estimators. In summary, *Ω*_T_ (Equation 12) contains both survivorship bias and truncation bias; *Ω*_TC_ contains only truncation bias (Equation 11) but this is because it is idealized (it can only be obtained from simulation data); *Ω*_o_ (Equation 9) contains only survivorship bias. In simulations, *Ω*_TC_ serves to illustrate the upper performance limit of *Ω*_T_ if the survivorship bias 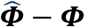 can be corrected.

The data and code used in this paper are available on a Github/Zenodo repository at https://github.com/EWTekwa/RichnessEstimator. The Matlab/R function, *RichnessEstsCov*, takes in a spatial abundance matrix and returns all estimators.

### Confidence Bounds

We produced 95% confidence bounds of each estimator from 2000 the full empirical datasets and 50 for each of 40 subsampled replicates within each scenario. We use non-parametric bootstrapping [35] because the richness distribution is unknown. Individual occurrences are not independent both within species and patches because of the intraspecific tendency to cluster [31,36]. In contrast, previous bootstrapping procedures for richness estimates assumed that sampled individuals are independent (as did the estimators themselves) [6], an assumption that is also used for rarefaction and extrapolation techniques that control for sampling effort when comparing communities [18]. To account for spatial and species dependencies of sampled individuals, for each bootstrap we first randomly sampled with replacement *k* transects (Figure 2). Then from this spatially randomized set we randomly sampled with replacement as many species as the point estimator predicts from the original dataset (eg. *Ω*_T_ or Chao1). All species are drawn from the observed set of *S* species but the bootstrap community now consists of a potentially larger number of pseudospecies. We used this bootstrap sample size instead of the raw *S*, because we know *S* underestimates the true community richness and bootstrapping is intended to recreate sampling variations from that community, not from the observed subset. We further used a mean bias-corrected percentile method to obtain the final confidence bounds. This involves first centering the bootstrap estimates so that the mean of the corrected bootstrap estimates matches the point estimate. Then the 2.5^th^ and 97.5^th^ percentiles of the corrected bootstrapped estimates serve as the 95% confidence bounds. For subsample schemes, the confidence bounds were plotted as means of the 2.5^th^ and 97.5^th^ quantiles across replicates, which showed average confidence bounds (not uncertainties in the richness point estimates across replicates). The bootstrapping function in Matlab/R is *BootRichnessEsts_all*.

**Figure 2.**
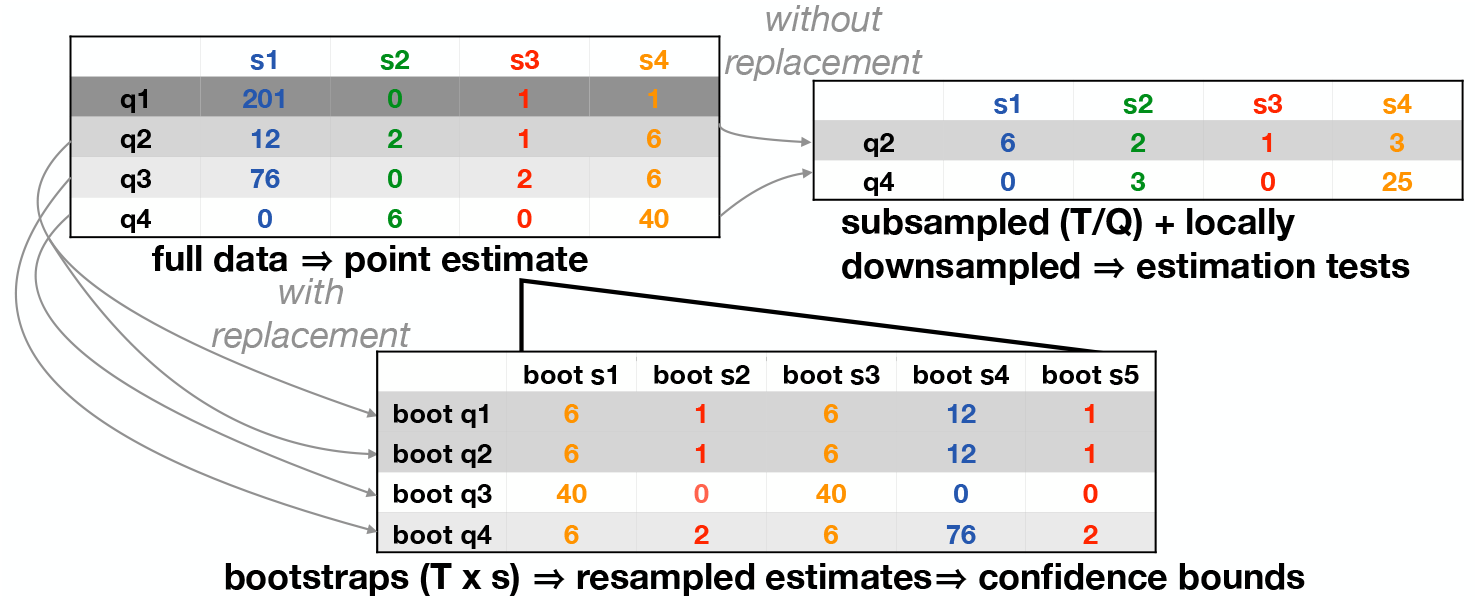
Bootstrapping, subsampling, and downsampling procedures. Any richness estimator produces a point estimate (eg. *Ω*_T_, here 5) from an original dataset (here 4 species *S* labelled by colours and 4 patches *q* labelled by grey shades). Each bootstrap first resamples with replacement patches from the original dataset, then resamples with replacments species for all resampled patches to create *Ω*_T_ pesudospecies. Multiple bootstrap estimates generate confidence bounds. Subsampling on empirial data tests how estimators perform with reduced spatial coverage, while downsapling is used to test how estimators perform with reduced individual coverage.

### Simulation Setup

To test the performance of asymptotic richness estimators, we simulated spatially partitioned communities under four scenarios that present different challenges that arise from imperfect local observation and spatial heterogeneities. For each simulation scenario, we created 2000 unique communities, each with 100 patches and 1 to 100 species. Communities in a scenario have different characteristics but converge on scenario mean values in abundance E[*n*], clustering E[*C*], and occupancy E[*P*], which simulate natural variations in spatial biodiversity patterns that we may want to discern. Further, each community was sampled at different efforts, converging on mean spatial sampling effort (patches sampled E[*k*]) and fraction of individuals observed E[*m*]. Thus, each of the 2000 communities represent a unique set of biological and sampling conditions.

Specifically, each simulated community’s mean fraction of individuals observed 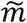 was ∼1/(1+(E[1/*m*]-1)*U*(0,1)) where *U* is a uniform random value. Each species in a community in turn had a 1*/m*_*i*_ value drawn from a normal distribution centered at the community mean 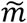 with a variance of 10 and a minimum of 1 (perfect observation). Each community’s mean abundance *ñ* was ∼E[*n*]*U*(0,1) and species *n*_*i*_ was drawn from a lognormal distribution with a standard deviation of 0.5. Each community’s mean clustering 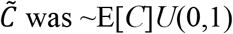, and species *C*_*i*_ was drawn from a lognormal distribution with a standard deviation of 1 and a minimum of 1 (no clustering). Each species’ *n*_*i*_ and *C*_*i*_ implied a *P*_*i*_ value according to the identity provided in [33]. We assigned individuals to each patch by drawing from a Poisson distribution with the mean being the species’ local density (Equation 1) multiplied by 1 if ∼*U*(0,1)<*P*_*i*_ and 0 otherwise. This led to true species abundance distributions that satisfied the scenario means E[*n*], E[*C*], and E[*P*]. Finally, subsampling biodiversity surveys were simulated by randomly picking *k*∈[2,10] patches in each community. Within each patch and for each species, individuals were observed by drawing from a Poisson distribution with the mean being the species’ local abundance times *m*. This implied sampling individuals with replacement within patches, such that locally observed abundance may be higher than what exists as *m* approaches 1, but even in this case the average observed individuals will converge to *n*. Sampling with replacement is likely in non-destructive field surveys when individuals move and their identities are not tracked; on the other hand the scheme approximates sampling without replacement when *m* is small. Our main results are from two simulation scenarios with E[*n*]=2: imperfect local observation (E[*m*]*<*1) (Figure 3A) and spatial heterogeneity (E[*C*]>1 and E[*P*]<1) (Figure 3B), each representing unique challenges to richness estimates. The *n* distributions for these scenarios are shown in Figure S1A-B. We also tested the estimators in scenarios with mixtures of imperfect local observation and spatial heterogeneity (Figure S2).

**Figure 3.**
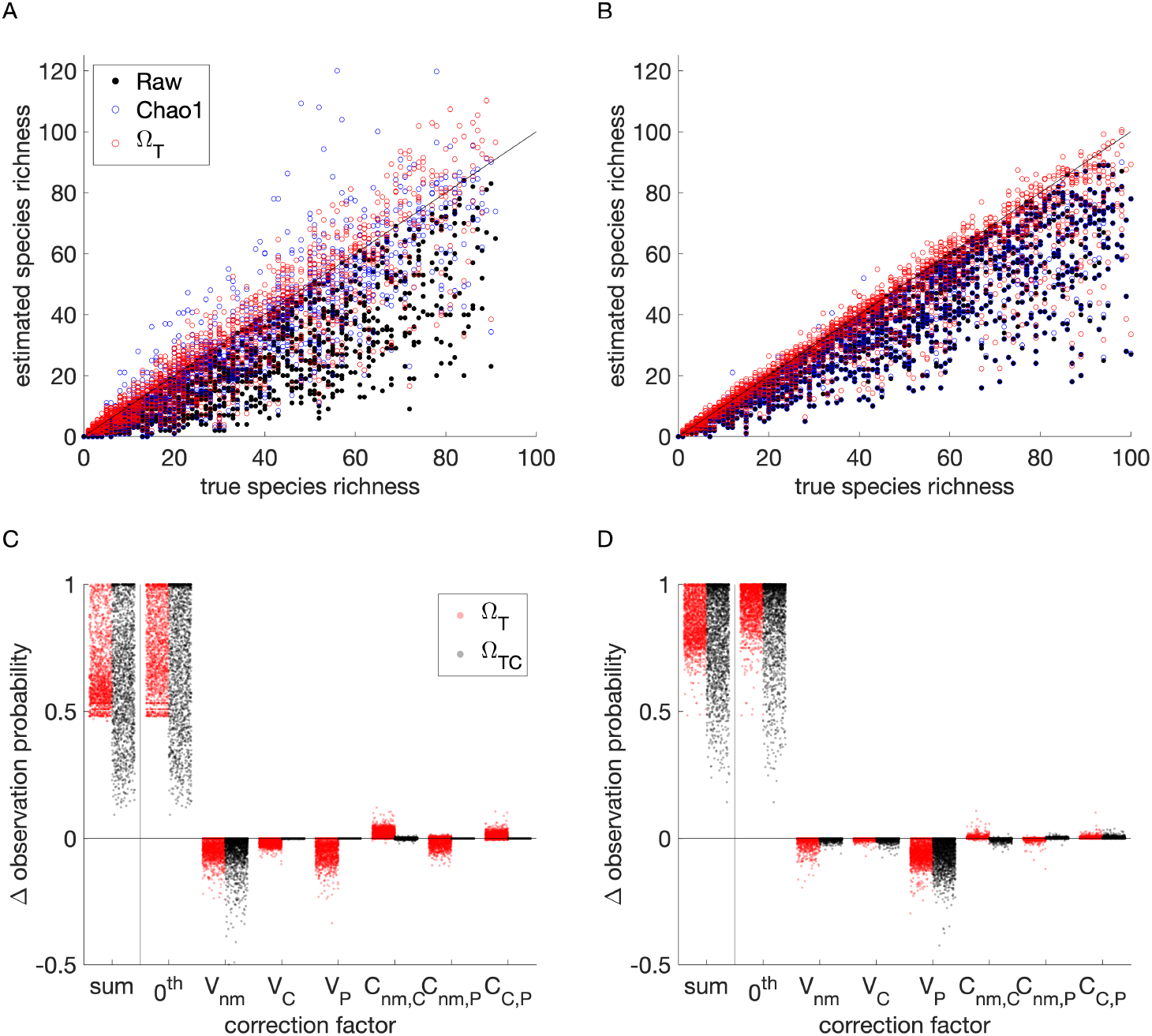
Simulated richness estimates. 2000 communities were populated with 1 to 100 species over 100 spatial patches, and 2 to 10 patches (*k*) were sampled for richness estimates, including raw (black dots), Chao1 (blue circles), and *Ω*_T_ (red circles). **A**. Imperfect local observation scenario, with mean parameter values of *m=*0.2, *n*=2, *C*=1, *P*=1, *k=*6. **B**. Spatial heterogeneity scenario, with mean parameter values of *m*=1, *n*=2, *C*=5, *P*=0.4, *k=*6. **C, D**. Corrections (Δ) to cross-species observation probability (with a baseline of one) by the proposed estimator (*Ω*_T_, red) and the corresponding idealized estimator where spatial abundance statistics were corrected for survivorship bias (*Ω*_TC_, grey), with the sum of correction factors (first column) broken into the individual factors (Equation 12). Each dot represents an estimate from the 2000 replicates, and ideally the *Ω*_T_ corrections would approach the *Ω*_TC_ corrections.

### Performance Evaluation

The evaluation of estimator’ performances in the simulations was conducted with six statistics that cover different aspects of bias, precision, and detection accuracy. A simple least-squares linear regression was performed on the estimated richness vs. true richness within each estimator. A slope less than 1 indicates that more diverse communities are more severely underestimated than less diverse communities, which is a prevalent bias among richness estimators (sometimes expressed as 1-slope [37]). In contrast to slope deviating from 1 as bias, precision is how close are multiple estimates given the same true richness; this can be measured by a low standard deviation (S.D.) in the regression slope. In addition, *R*^*2\**^ was computed for the 1:1 line, which describes the variance in estimated richness explained by true richness. A small *R*^*2\**^ indicates either underestimates or overestimates, with large errors capable of generating negative values. *R*^*2\**^ is an overall measure of the estimator’s accuracy. *R*^*2\**^ contrasts from the *R*^*2*^ of the true:estimate linear regression, the latter having been used to measure precision [37] but we believe the measure is confounded with bias. Even if an estimator always returns the same richness value for a given true richness, bias would reduce *R*^*2*^ below 1, thus we chose to use slope and S.D. in slope as our measures of bias and precision. Third, richness difference detections were checked between 2000 randomly picked community pairs that were different in richness by at most 2, 10, or 20 but were not identical. These tests provide more nuanced measures of detection accuracy relevant to uncovering spatial or temporal trends, which are major objectives of biodiversity studies. Detection accuracies above 0.5 is better than a coin-flip.

Low bias and high precision should improve accuracy, but there are many possible interpretations of accuracy. For instance, the detection accuracies we defined (richness difference detections in the paragraph above) refer to point estimates from the full dataset, whereas in practice we may look at whether spatial or temporal trend in richness change has a low average *p* value across bootstraps (when incorporating uncertainties). In this sense trend detection would be affected by an estimator’s precision more than pairwise comparison accuracy based on point estimates. We desire asymptotic richness estimators to both reduce bias and increase accuracy, but these components may exhibit tradeoffs given limited information [37]. Thus, the performance of an estimator cannot be summarized by a single measure; we therefore present all six measures introduced above to assess whether certain estimators rank higher in one aspect and lower in others.

Simulations from the worst data conditions led *Ω*_TC_ correction factors to estimate negative observation probabilities in up to 3% of the communities (Figure S2). These cases indicate conditions where variances and higher moments were unstable – even though *Ω*_TC_ had access to the true values of *mn, C*, and *P*, the data was still poor because *m* and *P* were small. Based on where outliers occurred, we set a threshold of 10% estimated observation probability below which *Ω*_T_ and *Ω*_TC_ switched to using only the 0^th^-order correction factor (first correction term in denominator of Equation 12), which cannot be negative and dominated among correction factors. This threshold did not affect *Ω*_T_ in the simulations but serves as a safeguard against unforeseen scenarios.

### Empirical Datasets

The first dataset that we will use to test estimators is the Barro Colorado Island tree census from Panama [26]. Start in 1982, every stem was identified by species every 5 years, for a total of 8 census. Over 300 species was observed across 1250 quadrats that form a continuous grid over 50 hectares. A negative temporal richness trend seems likely from the raw data. The census nature of the dataset allows us to conduct spatial subsampling and local downsampling experiments to test whether an estimator can recover the known true richness and whether the negative trend can be detected under poor sampling conditions.

The second dataset is an extensive seaweed diversity survey collected annually from 2012 to 2019 on the coast of British Columbia, Canada, and provide a protocol for obtaining bootstrapped confidence bounds. The dataset features relatively high standards in sampling effort and taxonomic identification for the marine realm [27]. Still, we expect uncertainty to persist because of imperfect sampling and spatial heterogeneity. We also conducted post-hoc subsampling experiments to explore how various scenarios of reduced sampling effort affect the estimators.

The survey tracked seaweed species and their abundance (estimated as percent cover) across steep gradients in abiotic stress associated with rapid changes in elevation and immersion time. The researchers established nine transects running parallel to shore at three intertidal elevations. Each year, we randomly selected ten quadrats on each transect and surveyed them for percent cover of all seaweed species. Between 73 and 82 species were observed each year, and no temporal trend can be detected from the raw data. Transects likely represent independent spatial samples of an underlying community, whereas quadrats were less spatially independent, so we aggregated quadrat data within transects for this analysis. A 10 × 10 grid was overlaid on the 0.5m x 0.5m quadrat, and any organism filling each square of this grid added 1 percent to the species’ total cover within the quadrat [38]. A species that was partially present in only one square of the grid was assigned a % cover of 0.5 (or trace). We therefore divided all percent cover data by 0.5 to obtain abundance.

Abundance within the context of sampling has no real biological meaning beyond being discrete; the abundance unit does not have to coincide with the individual organism, a concept that we note is open to biological debates and subject to evolution [39,40] but has no consequences for richness estimation. The only important feature of count data is that the observable abundance *mn*, not the biological *n*, approximates a Poisson process; clustering *C* and occupancy *P* are not affected by a change of unit and are thus insensitive to the individual definition (Equation 3 & Equation 5). The other consideration for converting percent cover to observed local abundance (*Cmn*) is that it is capped and thus not quite Poisson, but in practice recording abundances beyond this cap will likely not affect the estimated observation probability (eg. failure to observe, exp(-*Cmn*), is 4×10^−44^ for *Cmn* =100 in Equation 4).

In addition to obtaining point estimates of richness on both full datasets, we subsampled transects and quadrats in different combinations and produced replicated subsample estimates (plus bootstrap confidence bounds). For each subsampling replicate for the Barro Colorado Island tree census, we randomly pick a subset of quadrats (patches) without replacement and sampled these same quadrats every year. For each subsampling replicate for the British Columbia seaweed survey, we randomly pick transects and quadrats (without replacement) and sampled these same quadrats every year. Quadrat counts are summed to obtain transect data, so transects were treated as patches here. Each spatial effort level also receives a complement local downsampling scheme in which only 10% of individuals in each selected transect was observed. For each scheme, we replicate the subsampling 40 times since different transects and quadrats could be randomly picked. An ideal estimator would predict the same true richness regardless of spatial subsampling and local downsampling.

For each full dataset and subsampling/downsampling scheme, and for each subsampled replicate and bootstrap, we perform a linear regression of estimated richness on year or census. The slope of the regression is the detected temporal trend, the S.D. is the standard deviation in slope from the regression, and the *p*-value of the slope is a measure of how frequent are more extreme slopes expected given the null hypothesis of no trend. When the regression is performed on the point estimates (using the non-bootstrapped full data or subsampled data), the statistics are related to the detection accuracy that we measured in the simulation (presented in brackets in Figure 4 and Figure 5). For subsampled schemes, the point estimate regressions are performed across replicates and averaged to represent the expected statistics when sampling effort is lower than in reality. When the regression is performed on a bootstrapped dataset, the statistics include uncertainties in the estimates given the data, so the average *p*-value across bootstraps should be higher than for point estimates (presented as the main statistics in brackets in Figure 4 and Figure 5).

**Figure 4.**
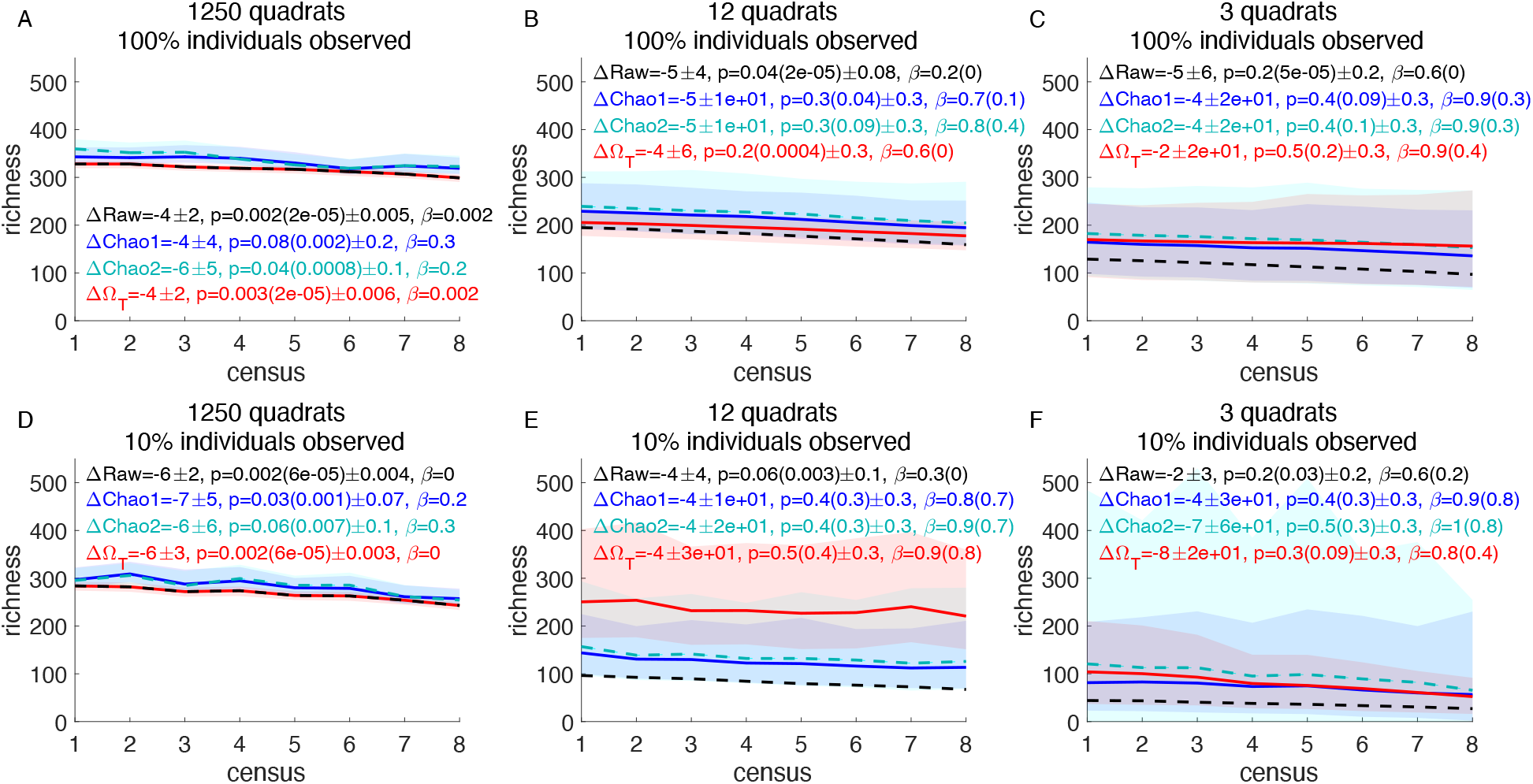
Richness estimates and bootstrap confidence bounds for the Barro Colorado Island tree census. For each year, raw (black dotted line), Chao1 (blue solid line), Chao2 (cyan dotted line), and *Ω*_T_ (red solid line) richness estimates were computed without **(A)** and with quadrat subsampling (40 times, each without replacement, same quadrats through years, **B, C**). Additionally we downsampled the portion of individuals observed in sampled patches **(D-F)**. Lines represent point estimates or mean point estimates across subsamples. 2000 total bootstraps were performed (50 for each subsampled dataset each census) for 95% confidence bounds. Shades represent upper (97.5^th^) and lower (2.5^th^) percentiles (or their means across subsamples). Displayed Δ are estimated changes in richness per census either from point estimates (bracketed statistics – ignoring observation and estimation errors) or from bootstrapped estimates. *β* is the observed false negative rate in detecting a temporal trend (frequency of *p*>0.05 across bootstraps and across subsamples) assuming there is a true trend (based on full census data).

**Figure 5.**
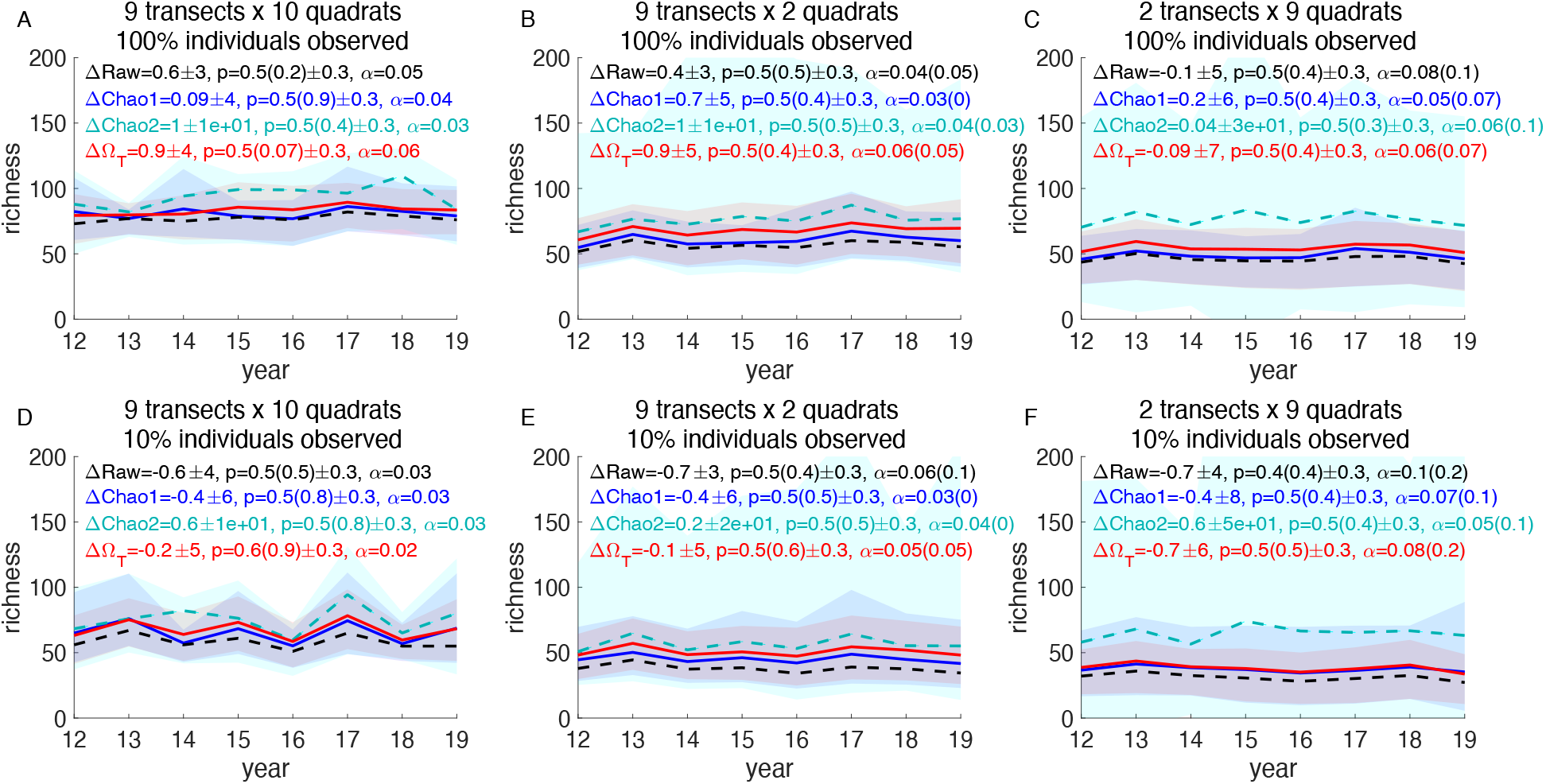
Richness estimates and bootstrap confidence bounds for the British Columbia seaweed survey. For each year, raw (black dotted line), Chao1 (blue solid line), Chao2 (cyan dotted line), and *Ω*_T_ (red solid line) richness estimates were computed without **(A, D)** and with transect and quadrat subsampling (40 times, each without replacement, **B, C, E, F**). Additionally we downsampled the portion of individuals observed in sampled patches in **(D-F)**. Transects were considered spatially independent samples of the community of species (*s*), and quadrat data were summed within transects. Lines represent point estimates or mean point estimates across subsamples. 2000 total bootstraps were performed (50 for each subsampled dataset each year) for 95% confidence bounds. Shades represent means of upper (97.5^th^) and lower (2.5^th^) percentiles across subsamples (or their means across subsamples). Displayed Δ are estimated changes in richness per census either from point estimates (bracketed statistics – ignoring observation and estimation errors) or from bootstrapped estimates. *α* is the observed false positive rate in detecting a temporal trend (frequency of p≤0.05 across bootstraps and across subsamples) assuming there is no true trend (based on full data).

For the Barro Coloardo Island tree census, we measure the false negative rate *β* (type II error) for detecting a temporal trend using each estimator (defined as *p*>0.05), assuming a trend is real. Per standard practice, *β*≤0.1 is considered an acceptable false negative rate; otherwise the study and estimator are statistically underpowered. For point estimates, *β* is obtained from the frequency of regressions across subsampled replicates whose slope *p*>0.05 (presented in brackets in Figure 4). These statistics do not exist for point estimates on the full dataset (because there is only one sample). For bootstrapped estimates, *β* is obtained from the mean frequency of *p*>0.05 across bootstraps (and all replicates under subsampling schemes). These bootstrapped *β* statistics are presented as the main statistics in Figure 4.

For the British Columbia seaweed survey, we measured the false positive rate *α* (type I error) for detecting a temporal trend using each estimator (defined as *p*≤0.05), assuming a trend does not exist. This null assumption is more tenuous than the negative trend assumption for the Barro Colorado dataset because the seaweed survey, like most marine studies, are not census and only covers a small fraction of the community habitat, so an estimated *α*>0.05 may either be a true false positive identification by an estimator or evidence for an actual temporal change. For point estimates, *α* is obtained from the frequency of regressions across subsampled replicates whose slope *p*≤0.05 (presented in brackets in Figure 5). These statistics do not exist for point estimates on the full dataset (because there is only one sample). For bootstrapped estimates, *α* is obtained from the mean frequency of *p*≤0.05 across bootstraps (and all replicates under subsampling schemes). These bootstrapped *α* statistics are presented as the main statistics in Figure 5.

## Results

### Theoretical Insights

Here we describe how sampled patches (*k*) and means and variances in observable abundance (*mn*), clustering (*C*), and occupancy (*P*) affect species observation and induce bias to the observed richness. First, we found that the effects of variable means 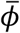 (across species and patches) were captured by the 0^th^-order correction factor 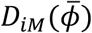 in the Taylor expansion of the richness estimator (Equation 11), which is the single-species observation probability (Equation 7) computed at variables’ mean values (as if all species in the community share the same characteristics). Equation 7 states that observation probability increases with means of *k, n, m, C*, and *P*.

Across species variance (var(*ϕ*)) effects depend on the signs of the second-order partial derivatives (Equation 11). The analytical expressions are long but always negative (see numerical outcomes in Figure 3C-D). Therefore, variances always decrease the overall observation probability.

These results clarify two main ways in which non-richness biodiversity components induce bias to observed richness. Clustering and occupancy are related to spatial turnover and dissimilarity, yet their mean values affect species observation (Equation 7). Variance in observable abundance, which is likely inversely proportional to community evenness (assuming *m* is not anti-correlated with *n*), reduces observed richness even though true richness should not be affected by evenness. These effects illustrate the need for biodiversity metrics to be bias-corrected if they are to represent their definitions.

### Simulation Tests

We tested the proposed estimators (*Ω*_o_: Equation 9 and *Ω*_T_: Equation 11) on a simulated dataset of species abundance across a heterogeneous landscape. We compared *Ω*_o_ and *Ω*_T_ to other leading estimators including: the unbiased Chao1 [6,22], Chao2 [28], the Abundance-based coverage estimator (ACE) [41], Jackknife1 for abundance data, and Jackknife2 for incidence data [42]. We also tested the idealized version of *Ω*_T_ that is only opertional in simulations where the true spatial statistics are known for all species to establish theoretical limits of *Ω*-type estimators (*Ω*_TC_: Equation 11). For incidence-based estimators including Chao2 and Jackknife2, abundance within patches were converted to presence/absence. The six performance statistics in Table 1 are described in the Methods: Performance Evaluation section.

**Table 1.** Simulation results. Performance was measured as the slope of estimated richness when regressed on true richness (downward bias when <1), standard deviation in regression slope (precision), *R*^*2\**^ of the 1:1 line (overall accuracy), and pairwise difference detection accuracy when randomly picked communities (out of 2000) were within a richness difference of 2, 10, or 20 (detection accuracy). For each scenario **(A-D)** and statistics, ranks higher than raw are indicated by dark shades (with black indicating top ranks); ranks lower than raw are unshaded.

| Estimator | A. Imperfect local observation<br>($m=0.2, n=2, C=1, P=1, k=6$) | | | B. Spatial heterogeneity<br>($m=1, n=2, C=5, P=0.4, k=6$) | | |
| --- | --- | --- | --- | --- | --- | --- |
| | Slope±S.D. | 1:1<br>$R^{2*}$ | Detection<br>($\leq \Delta 2, \leq \Delta 10, \leq \Delta 20$) | Slope±S.D. | 1:1<br>$R^{2*}$ | Detection<br>($\leq \Delta 2, \leq \Delta 10, \leq \Delta 20$) |
| $\Omega_{TC}$ | 0.93 ± 0.26 | 0.95 | 0.59, 0.79, 0.88 | 1.0 ± 0.15 | 0.98 | 0.64, 0.84, 0.91 |
| $\Omega_o$ | 0.92 ± 0.43 | 0.88 | 0.59, 0.73, 0.80 | 0.87 ± 0.31 | 0.89 | 0.61, 0.76, 0.83 |
| $\Omega_T$ | 0.95 ± 0.46 | 0.87 | 0.58, 0.73, 0.79 | 0.86 ± 0.30 | 0.89 | 0.62, 0.75, 0.83 |
| Chao1 | 0.93 ± 0.43 | 0.80 | 0.56, 0.70, 0.78 | 0.74 ± 0.30 | 0.76 | 0.58, 0.71, 0.77 |
| Chao2 | 1.0 ± 0.58 | 0.80 | 0.56, 0.70, 0.78 | 0.97 ± 0.37 | 0.88 | 0.60, 0.76, 0.84 |
| ACE | 1.0 ± 0.46 | 0.79 | 0.56, 0.70, 0.79 | 0.74 ± 0.30 | 0.76 | 0.58, 0.72, 0.78 |
| Jackknife1 | 1.0 ± 0.33 | 0.89 | 0.54, 0.68, 0.79 | 0.75 ± 0.31 | 0.76 | 0.56, 0.69, 0.76 |
| Jackknife2 | 1.1 ± 0.47 | 0.88 | 0.52, 0.63, 0.72 | 1.1 ± 0.35 | 0.88 | 0.55, 0.72, 0.81 |
| Raw | 0.65 ± 0.41 | 0.64 | 0.52, 0.65, 0.73 | 0.73 ± 0.32 | 0.75 | 0.57, 0.70, 0.76 |

| Estimator | C. Mixed conditions<br>($m=0.2, n=2, C=5, P=0.4, k=6$) | | | D. Mixed poor conditions<br>($m=0.1, n=2, C=8, P=0.3, k=6$) | | |
| --- | --- | --- | --- | --- | --- | --- |
| | Slope±S.D. | 1:1<br>$R^{2*}$ | Detection<br>($\leq \Delta 2, \leq \Delta 10, \leq \Delta 20$) | Slope±S.D. | 1:1<br>$R^{2*}$ | Detection<br>($\leq \Delta 2, \leq \Delta 10, \leq \Delta 20$) |
| $\Omega_{TC}$ | 1.2 ± 0.76 | 0.51 | 0.55, 0.70, 0.78 | 1.1±1.1 | 0.01 | 0.53, 0.60, 0.69 |
| $\Omega_o$ | 0.75 ± 0.48 | 0.66 | 0.54, 0.63, 0.71 | 0.52 ± 0.51 | 0.04 | 0.53, 0.55, 0.64 |
| $\Omega_T$ | 0.78 ± 0.51 | 0.68 | 0.54, 0.64, 0.71 | 0.55 ± 0.55 | 0.09 | 0.53, 0.54, 0.64 |
| Chao1 | 0.67 ± 0.46 | 0.49 | 0.54, 0.62, 0.68 | 0.53 ± 0.54 | -0.06 | 0.50, 0.53, 0.62 |
| Chao2 | 1.0 ± 0.79 | 0.51 | 0.52, 0.64, 0.72 | 0.97 ± 1.0 | -0.08 | 0.51, 0.54, 0.63 |
| ACE | 0.69 ± 0.53 | 0.51 | 0.54, 0.62, 0.69 | 0.60 ± 0.65 | 0.04 | 0.51, 0.53, 0.61 |
| Jackknife1 | 0.74 ± 0.44 | 0.64 | 0.51, 0.61, 0.69 | 0.55 ± 0.46 | 0.14 | 0.49, 0.53, 0.61 |
| Jackknife2 | 1.0 ± 0.54 | 0.79 | 0.50, 0.60, 0.70 | 0.74 ± 0.73 | 0.39 | 0.46, 0.46, 0.56 |
| Raw | 0.51 ± 0.39 | 0.22 | 0.52, 0.58, 0.67 | 0.33± 0.37 | -0.62 | 0.49, 0.50, 0.61 |

The simulations show that our proposed *Ω*_o_ and *Ω*_T_ estimators were either the best or close to the best across observation and spatial heterogeneity scenarios in terms of detection accuracy (across all true difference levels of 2, 10, or 20), while their biases (true:estimated richness slope) were generally lower than other abundance-based estimators but higher than incidence-based estimators (Figure 3, Figure S2, Table 1). All estimators eroded precision when compared to raw, but *Ω*_o_ and *Ω*_T_ had the highest precision after raw (in term of standard deviation of slope estimate). Incidence-based estimators performed better under spatial heterogeneity [37] but worse under imperfect local observations. In particular, Chao2 and Jackknife2 for incidence data contained the least bias as a function of true richness (in term of regression slope) but performed poorly in detecting small differences under poor conditions (Table 1 C&D). Jackknife2 consistently ranked near the top along with *Ω*_o_ and *Ω*_T_ in term of overall accuracy (1:1 *R*^*2\**^), but Jackknife2 detected actual differences much less accurately. All estimators other than Chao1, *Ω*_o_, and *Ω*_T_ were near-worst performers under some scenarios and measures so are prone to catastrophic failures when the underlying community characteristics are unknown. Between the three most robust estimators, *Ω*_T_ always out-ranked Chao1 and either out-ranked or equaled *Ω*_o_ across all scenarios and measures. In particular, under the poorest scenario only *Ω*_o_ and *Ω*_T_ were better than coin flips in detecting richness differences were ≤2 (scenario D in Table 1).

Between *Ω*_o_ and *Ω*_T_, we found that *Ω*_T_ exhibited less bias overall while losing little in precision and detection accuracy, therefore we will focus on *Ω*_T_ as the main product of our paper (Equation 12). The proposed *Ω*_T_ gained 2-8% in detection accuracy compared to using raw observed richness across scenarios. The idealized *Ω*_TC_ (only operational in simulations) performed best overall, further improving difference detection accuracy (up to 15% higher than raw) while almost eliminating bias (slope close to one, grey rows in Table 1).

The room for improvement to *Ω*_T_ comes from survivorship bias in raw measurements of *mn, C*, and *P*. This bias can be observed in the differences between the *Ω*_T_ and *Ω*_TC_ correction factors in each simulation scenario (Figure 3C-D). Across both test sets, observed mean and variance in *mn* were biased upward because rare species were missed and observation noise added to differences (Figure S1C, E). The upward bias in mean *mn* overestimated observation probability, but the upward bias in *mn* variance underestimated observation probability (Figure 3C-D). For the imperfect local observation scenario, mean observed *P* was biased downward, and mean observed *C* and variances in *C* and *P* were all biased upward (Figure S1D). The downward bias in mean *P* and upward biases in variances in *C* and *P* led to underestimates in observation probability (Figure 3C), while the upward biases in mean *C* overestimated observation probability. For the scenario with spatial heterogeneity, the biases in means and variances of *C* and *P* were the exact opposite of those caused by low fractions of individuals observed (Figure 3D, Figure S1F). The opposing effects of biases across variables partly cancelled each other, reducing net bias in the *Ω*_T_ richness estimate and causing the net correction to approach that of the idealized estimator *Ω*_TC_ (Figure 3C-D). The covariance correction factors were generally smaller than the other factors and similar in *Ω*_0_ and *Ω*_T_. In comparison, other estimators depended only on the observation of rare species and did not benefit from opposing spatial or variational biases [6]. However, the net bias in our method depended on the relative magnitudes of these survivorship biases, which explains why unbiased versions of the variables (inaccessible in real datasets) improved the proposed method (up to *Ω*_TC_). Consistent directions in the survivorship biases associated with different biotic and abiotic sources suggest that corrections are possible to some extent; we identify these as future research objectives.

Next we examine how well the proposed estimators perform in reducing bias and detecting differences in real applications where sampling is incomplete.

### Barro Colorado Island Tree Census

We first computed the raw observed richness, Chao1, Chao2, and the proposed *Ω*_T_ point estimates for each year using the full census data. For *Ω*_T_, observed spatial statistics used to make corrections are shown in Figure S3. Based on the raw data, a negative trend of four species lost per census (per 5 years) was observed (Figure 4A, *p*=2×10^−5^ from a least-squares linear regression). Bootstrapped regressions on the raw data (containing possible variations of the true community represented by the census plot) continued to support a negative temporal trend (mean *p*=0.002). We therefore recorded the bootstrapped false negative rate *β* for all estimators across subsampling schemes assuming a true negative trend (assuming a significance level *α*=0.05 and a standard acceptable false negative rate being *β*<0.1). Applications to bootstrapped datasets reveal that Chao1 and Chao2 overcorrected and obscured the temporal trend, as seen by increased *p* values and *β*>0.1. In contrast, *Ω*_T_ determined no correction was necessary given the data quality and retained the original temporal trend, *p* values, and *β*.

Next we examined how the estimators perform when only 12 or 3 quadrats out of 1250 were sampled through the census years. Across 40 replications for each of the spatial subsampling schemes, all estimators except for the raw data at 12 sampled quadrats indicate no evidence of temporal trend and high false negative rates when bootstrapped uncertainty was included (high main *p* and *β* values in Figure 4B, C). However, when only the point estimates are considered (excludes uncertainty in richness estimates), both raw and *Ω*_T_ consistently indicated a decreasing temporal trend at 12 sampled quadrats, and raw continued to do so at 3 sampled quadrats (low bracketed *p* and *β* values in Figure 4B, C). When spatial subsampling was coupled with local downsampling of individuals observed, all except raw failed to detect a temporal trend (Figure 4E, F), although *Ω*_T_ came close to recovering the trend at 3 sampled quadrats based on point estimates (*p*=0.09 in Figure 4F). While a generally lower *β* for *Ω*_T_ when compared to Chao1 and Chao2 across sampling schemes indicates a greater derived statistical power (lower data requirement to infer the true temporal trend), regressions based on raw data consistently indicated the lowest *β*. In this dataset raw data correctly identified the trend even with subsampling and downsampling, which appears to oppose the simulation conclusion that raw data is worst at detection. *Ω*_T_’s correction at the intermediate spatial sample size was likely suppressed by the non-independence of quadrat samples, especially without downsampling (effective *k*_*i*_ in Equation 7 is likely smaller than the number of quadrats sampled). We will revisit the apparent contradiction in detection accuracy in the Discussion.

Overall, *Ω*_T_ exhibited similar or less downward bias compared to other estimators (including raw) under all subsampling and downsampling, recovering close to the raw absolute richness from the full dataset when only 12 quadrats and 10% of individuals were sampled (Figure 4E). Precision decreased in this case, but detection accuracy was no worse than other estimators except when compared to raw. In all subsampled and downsampled schemes, only Chao2 (Figure 4D and F) and *Ω*_T_ (Figure 4E) had confidence bounds that sometimes covered the raw richness from the full dataset. These results are consistent with the simulation analyses, which revealed Chao2 and *Ω*_T_ should be less biased than Chao1.

### British Columbia Seaweed Survey

In this marine seaweed diversity dataset, the raw observed richness across years indicated a marginal signal of temporal change but was inconclusive (average *p*=0.5 across bootstraps or 0.2 for point estimate, *α*=0.05 fraction of times bootstrapped regression showed *p*≤0.05, Figure 5A). For *Ω*_T_, observed spatial statistics used to make corrections are shown in Figure S4. Chao1 and Chao2 indicated no temporal trend, while *Ω*_T_ indicated marginal support for increasing richness based on point estimates (*p*=0.07, *α*=0.06 across bootstraps). Given these statistics, the true temporal trend is likely weak and the null hypothesis of no change prevails. *Ω*_T_ indicated that between 3 to 9 seaweed species remain unobserved in the survey throughout the years, while Chao1 indicated none or few species unobserved. Chao2 made larger upward corrections that were uneven across years. When the full spatial sample was locally downsampled, all estimators indicated no temporal trends (Figure 5D).

Under the spatial subsampling scheme of 9 transects with 2 quadrats each, all estimators including raw showed no evidence of temporal trends (Figure 5B). Confidence bounds for Chao2 and *Ω*_T_ covered the raw richness from the full dataset, but the upper bound for Chao2 was well beyond the probable truth (>200 in some years). Under the subsampling scheme of 2 transects with 9 quadrats each, raw and Chao2 were more prone to false positives than Chao1 and *Ω*_T_ (*α* in Figure 5C). Only Chao2’s confidence bounds covered the full dataset’s raw richness but again the upper bound appears too high. With the addition of local downsampling on top of spatial subsampling, raw data continued to show the highest false positive rates (*α* in Figure 5E-F).

Overall, in this dataset Chao2 exhibited the least bias but the lowest precision, while *Ω*_T_ was the most precise, less biased than Chao1, and potentially detected a stronger signal of temporal trend in the full dataset.

## Discussion

The proposed *Ω* richness estimators for species richness achieve a superior balance of low bias while retaining precision and accuracy in detecting richness differences compared to abundance-based estimators including Chao1 [6], ACE [41], and Jackknife1 [22]. In particular, our second-order Taylor-expansion estimator *Ω*_T_ (Equation 12) accounts for the mean and variance effects of patches sampled (sample size), fractions of individuals observed within each species (catchability), abundance (rarity), clustering (spatial variance), and occupancy, all readily computed from spatial abundance data and without the need to independently assess survey effectiveness. *Ω*_T_ harnesses information already present in common surveys, thus it is no more data-hungry than other methods we assessed. The proposed method can be used to study spatial and temporal trends [17] as well as their relationships with other environmental covariates, without the confounding effects of these covariates on richness estimates (i.e. richness and other covariates may spuriously influence each other through endogeneities introduced if richness is first estimated with the aid of those covariates) [20,21].

There remains a downward bias (increasing underestimation when true richness increases) to all richness estimators we examined with simulations including *Ω*_T_, with the exception of the incidence-based Chao2 [28] and Jackknife2 [22], which surprisingly had the minimal bias when looking across all replicates despite ignoring abundances (they only relied on spatial occupancy). However, these incidence-based estimators were poor at resolving small differences between communities under data-poor conditions. In contrast, across the different scenarios that simulations datasets represent, *Ω*_T_ detected trends correctly more often than no correction and any other estimators under any condition while maintaining the lowest bias among abundance-based estimators. Nevertheless, *Ω*_T_ contains survivorship bias [34] because observed species tend to have higher abundance, lower clustering, and higher occupancy than unobserved species. If survivorship bias can be corrected (as represented by *Ω*_TC_ in simulations) and this may very well be possible with further research because the bias directions are known the proposed approach would come very close to eliminating bias while increasing detection accuracy drastically. Like Chao1, *Ω*_T_ is best understood as a lower-bound richness estimator but for slightly different reasons. Chao1 was explicitly derived as a lower-bound estimator regardless of moment truncations [6], whereas the downward bias for *Ω*_T_ came from survivorship bias. In addition, non-independence of spatial samples (*k*) adds downward bias to all *Ω* estimators by inflating inferred observation probabilities; a careful treatment of effective sample size in the context of these estimators would help improve performance further.

Interestingly, the second-order Taylor-expansion estimator *Ω*_T_ outperformed the exact estimator *Ω*_o_ derived directly from the inferred observation probabilities across observed species. In other words, the Taylor series truncation provided a better description of the underlying community than assuming that the observed species were the best representations of the hidden community. This is surprising because *Ω*_o_ contains only survivorship bias (since the spatial abundance statistics are only available for observed species), whereas *Ω*_T_ contained both survivorship bias and truncation error (from excluding statistical moments beyond the second order). While it is well-known that higher moments are individually difficult to estimate and would thus introduce instability to the estimator, it remains to be mathematically explained why the sum of all higher moments up to infinite order, which are theoretically contained in *Ω*_o_, put *Ω*_o_ at a slight disadvantage when compared to *Ω*_T_. However, the theme of multiple opposing biases also operates among the six different correction factors within the second-order Taylor approximation of species observation probability in *Ω*_T_. The magnitudes and directions of these biases are sensitive to underlying non-spatial and spatial heterogeneities in the community, which always partially cancel each other, which helps explain why the sum outperforms existing estimators.

Empirical applications of the estimators to the Barro Colorado Island tree census and the British Columbia seaweed survey revealed some expected patterns and other surprising results. Using spatial subsampling and local downsampling of individuals, richness estimates and bootstrapped confidence bounds from Chao1, Chao2, and *Ω*_T_ were compared to those obtained using the full datasets. These comparisons showed that *Ω*_T_ and Chao2 were less biased than Chao1 and raw; under different conditions either *Ω*_T_ or Chao2 was the least biased (had estimates from subsampled data that were closest to richness observed in the full dataset). *Ω*_T_ was more precise than Chao1 and Chao2, the latter being very imprecise with poor data. Application to the Barro Colorado Island data showed that *Ω*_T_ was superior to Chao1 and Chao2 in detecting the presumably true negative temporal trend across census years either with the full data or with subsampled and downsampled data (lower false negative rate *β*); in other words *Ω*_T_ gains more statistical power than other corrective estimators and thus requires less data despite the estimator’s apparent complexity. However, the raw data was surprisingly more consistent than *Ω*_T_ in detecting a negative trend under poor data conditions, which contradicts our simulation findings that raw data should be the worst at detection. But we should keep in mind this is only one counterexample of possible communities; experiments with different datasets containing clear temporal or spatial trends will be necessary to answer whether our simulation conclusions hold on average. The related question of false positive rate associated with estimators was more clearly answered with the British Columbia seaweed survey, where the true temporal trend is likely constant across years. There the raw data was most prone to falsely detecting a trend under subsampling and downsampling. A caveat to the assumption of no true change is that *Ω*_T_ was the only estimator that generated marginal support for a slight richness increase in the full dataset. Overall, the empirical applications demonstrated some superior qualities of *Ω*_T_ that parallel the simulation results, but our uncertainty about the true states of the communities, as well as the paucity of different communities examined, leave room for further empirical and simulation assessments.

We note that *Ω*_T_ is not restricted to counting species and individuals – the number of any taxonomic unit or non-biological class can be estimated instead of species (e.g., order, family, functional, or morphological richness, alphabets, words, etc.), and any discrete abundance unit can replace the biological individual counts. The method partly relies on individual units being sampled within-patch through a Poisson process, which is likely satisfied even by survey methods that appear to carry continuous values. Beyond this, the method should even handle spatial incidence data. With local observable abundance *C*_*i*_*m*_*i*_*n*_*i*_ fixed to a sufficiently high value for observed species (set to >10 when preparing the data), the observation probability per occupied patch (Equation 4) is effectively one, meaning that the only effect on *Ω*_T_ is equating the observation probability per patch *D*_*ix*_ with occupancy *P*_*i*_ (Equation 6). This simple extension illustrates the potential for the current approach to handle a variety of data qualities including surveys designed for biodiversity assessment, unstructured observational data, and mixed data meta-analyses. We leave it for future research to quantify and improve the breadth and performance of *Ω*-type richness estimators across different data types. One can further examine whether rarefaction and extrapolation techniques that control for sample effort or coverage [18,23] can improve difference detection accuracy when it is not a priority to estimate absolute richness (i.e. to reduce bias). This would not be an easy endeavour however, as the spatially heterogeneous setting that our estimator explicitly confronts also poses unaddressed problems on how to properly measure sample coverage.

While we have offered an improved asymptotic richness estimator that already performs well in balancing bias, precision, and accuracy, we identified many more open questions and precise research venues that may be large undertakings on their own. These complications reveal the inherent difficulties in estimating species richness and detecting biodiversity change, but also point to exciting possibilities that come with an approach that deviates completely from past methods. Ultimately, any richness estimator will be sensitive to whether the observed species are representative of the underlying community; in fact, the meaning of the underlying community in terms of taxonomic and geographic inclusion will be defined by the organisms that we choose to observe and record in the first place. Thus, even when armed with an ideal richness estimator, biological and field-methodological considerations are indispensable to improving the accuracy of estimated richness and interpreting its meaning across different communities.

## Acknowledgement

We thank the Hakai Biodiversity Synthesis group for discussions. ET and MW were funded by the Tula Foundation. ET was additionally funded by a Mitacs Accelerate Internship. PM was funded by a NSERC Discovery Grant (RGPIN 2019-06240).

