## Supplementary material for "Theory and application of an improved species richness estimator": Figure S

### Electronic Supplementary Material for


### Supplementary Figures

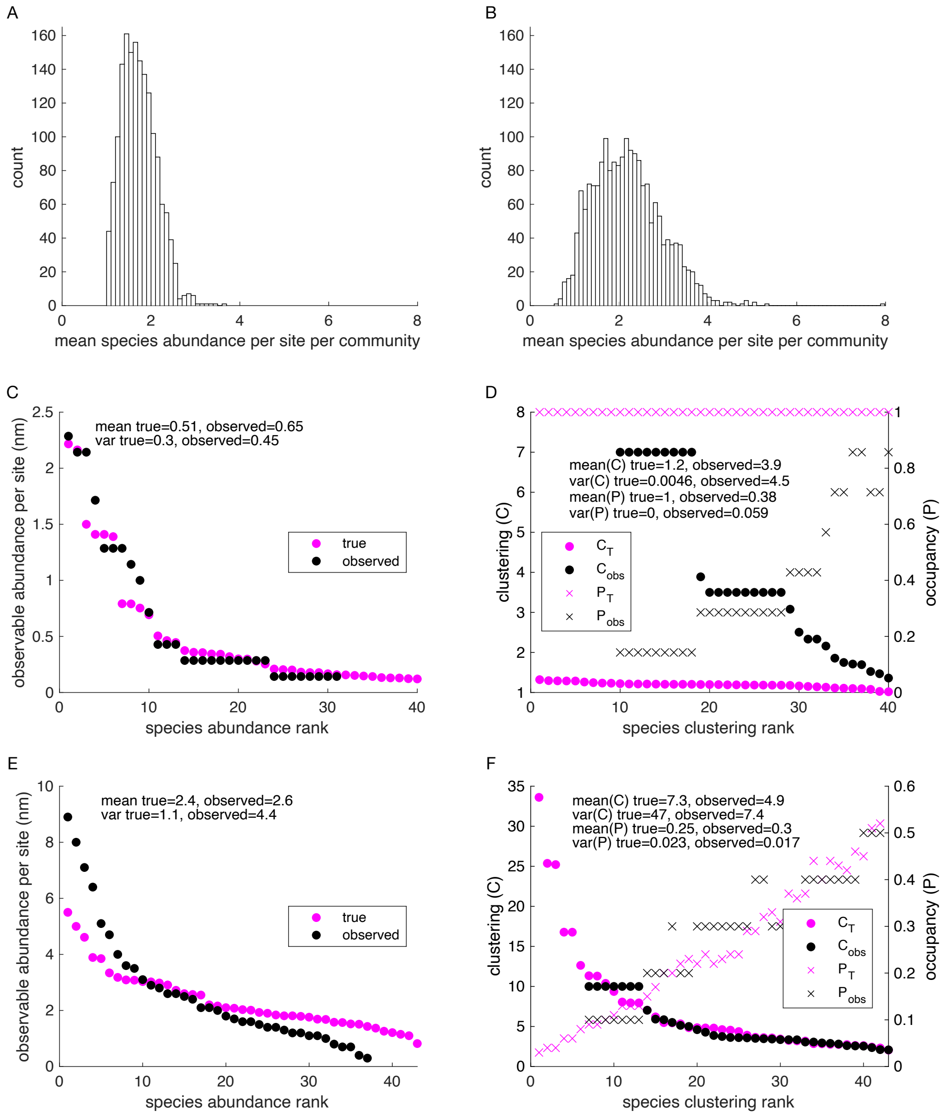


**Figure S1. Simulation measurements. A-B.** True mean species abundances per site among 2000 simulated communities (corresponding to Figure 2A and B parameters respectively). **C & E.** True (magenta) versus observed (black) observable abundance per site (*mn*) according to abundance rank. **D & F.** True (magenta) versus observed (black) clustering (*C*: dots) and occupancy (*P*: crosses) according to clustering rank. **C-D.** One simulated community corresponding to imperfect local observation (**Error! Reference source not found.**A parameters). **E-F.** One simulated community corresponding to high spatial heterogeneity (**Error! Reference source not found.**B parameters).

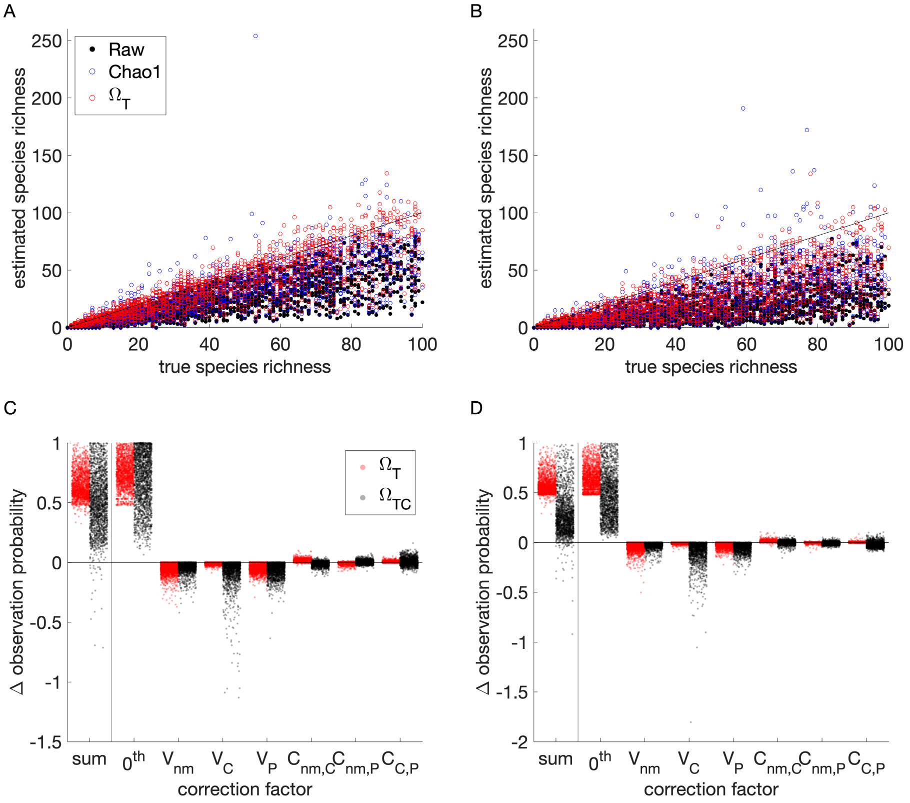


**Figure S2. Simulated richness estimates under mixed conditions. A.** Imperfect local observation and spatial heterogeneity scenario. Average parameter values are $m$=0.2, $n$=2, $C$=5, $P$=0.4, $k$=6. **B.** Poor local observation and highly spatial heterogeneity scenario. Average parameter values are $m$=0.1, $n$=2, $C$=8, $P$=0.3, $k$=6. Graphic descriptions are the same as for Figure 2. Note negative sum Δ observation probabilities for $\Omega_{TC}$ in **C** and **D** indicated situations where second-order truncation is unstable; in these cases and all other cases where sum Δ observation probabilities<0.1 only the $0^{th}$-order correction factor was used.

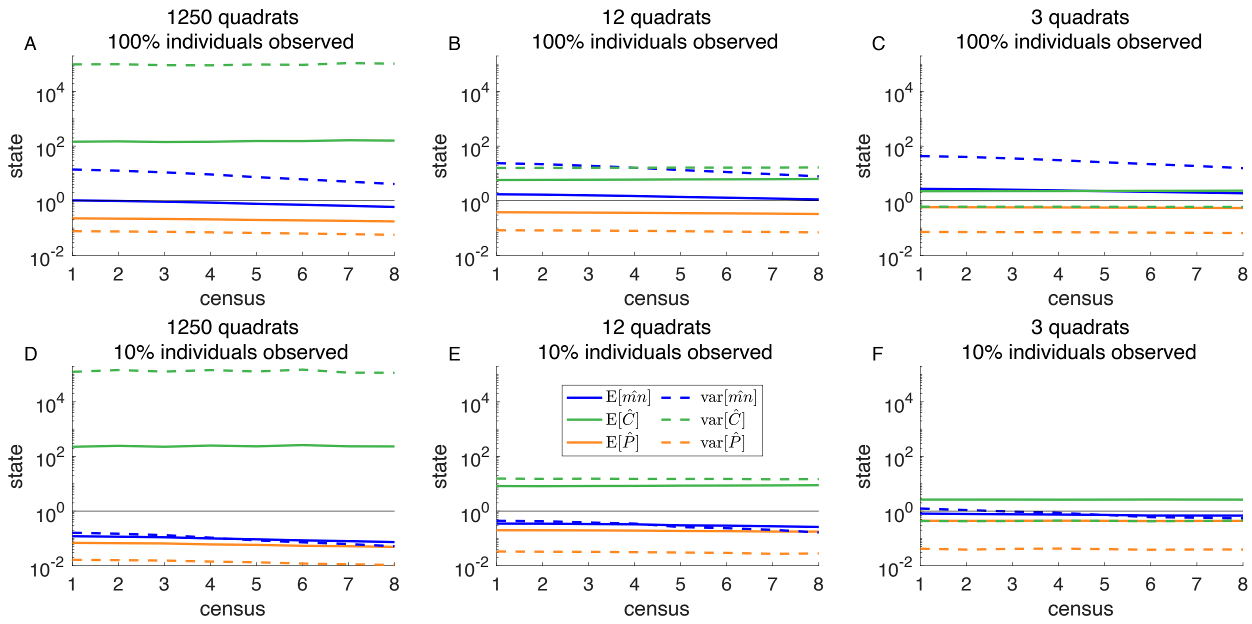


**Figure S3. Spatial statistics from the Barro Colorado Island tree census.** The observed states (means: solid lines, and variances: dotted lines) of abundance *mn* (blue), clustering *C* (green), and occupancy *P* (orange) among observed species. Reference black line has the value of one. **A.** Full dataset. **B-C.** Spatially subsampled data. **D-F.** Locally downsampled data. For **B-C** and **E-F**, states are averaged over 40 replicates.

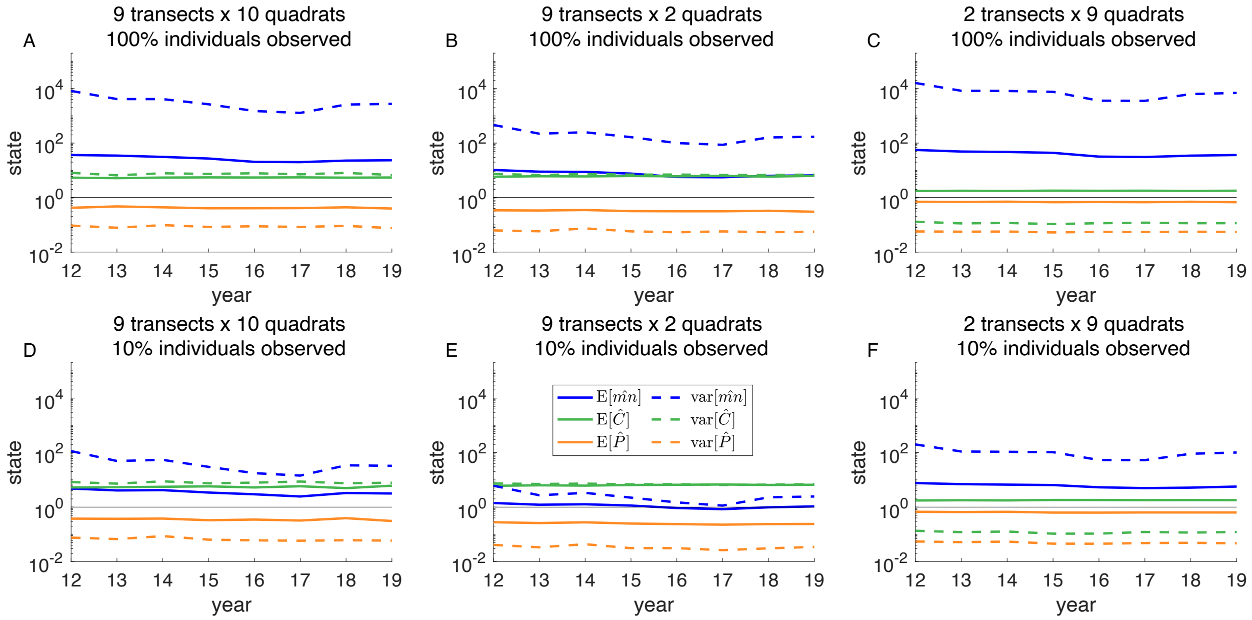


**Figure S4. Spatial statistics from the British Columbia seaweed survey.** The observed states (means: solid lines, and variances: dotted lines) of abundance *mn* (blue), clustering *C* (green), and occupancy *P* (orange) among observed species. Reference black line has the value of one. **A.** Full dataset. **B-C.** Spatially subsampled data. **D-F.** Locally downsampled data. For **B-C** and **E-F**, states are averaged over 40 replicates.
